## Supplementary Materials for "Assessing emergence risk of double-resistant and triple-resistant genotypes of *Plasmodium falciparum*"

Supplementary Information

#### 1 Specific MDR risks when MFT generates more risk than a cycling strategy

Figures 4 and 5 of the main text show that across the five maximally-resistant MDR genotypes we defined, MDR risk – as defined by the area under the genotype-frequency curve (AUC) – is lowest under MFT policies. However, this does not mean that MFT generates the lowest frequencies of each individual genotype, just the lowest sum of frequencies across all genotypes. Supplementary Figures 1 to 22 show the genotype frequencies in all epidemiological scenarios investigated here, and the underlined parts of each caption highlight the comparisons for which MFT is associated with higher MDR risk than one or both cycling policies.

In some scenarios, MDR generated slightly higher risk for the ASAQ double-resistant, the reason being that the simulations start with all genotypes carrying some resistance to AQ (as is true in reality). Therefore, with MFT using AQ for 33% of cases, 5-year cycling using AQ for 25% of cases, and adaptive cycling using AQ for about 10% to 15% of cases (depending on each individual simulation and when drug switches occur), MFT clearly puts the most selection pressure on AQ-resistant genotypes. Note that this is for a non-adaptive MFT approach that naively deploys three ACTs in equal amounts for 20 years without adjusting to changing levels of drug resistance. In Supplementary Table 1, we list all 8 of these scenarios and show that a simple change to a 50/50 MFT deployment of AL and DHA-PPQ results in substantial reductions in MFT's risk of driving an ASAQ double-resistant to high frequencies.

Note that the AUC numbers are not adjusted for prevalence. In other words, these numbers are in units of frequency-days not infection-days. This means that for some of the scenarios (e.g. 0.1% prevalence and 60% treatment coverage), an AUC value of 200 corresponds to 200 days of a genotype being fixed (frequency = 1.0), but this genotype could be fixed in a group of 50 parasite-positive individuals. For a scenario with MDR importation, some of these individuals may be recent imports meaning that their presence was not influenced by the drug policy in place.

Finally, a low absolute AUC value does correspond to low risk of an MDR genotype being generated during a 20-year period. AUC values in Supplementary Table 1 that are in the single digits correspond to an equivalent risk of a single-digit number of days (out of a twenty year period) that a genotype is fixed in the population and thus guaranteed to be the one transmitted onward if a parasite-positive individual is bitten by a mosquito.

#### 2 Definition of median simulation for Figures 6 and 7

To construct a mutation-flow diagram a representative simulation must be chosen out of the 100 simulated for each scenario. To do this, the median frequency of each genotype is calculated, by month, for the entire 20-year duration of the simulation. This gives $12 \times 20 \times 5 = 1200$ genotype-frequency data points for the five maximally-resistant genotypes. The simulation with the minimum absolute distance (summed over these 1200 data points) to the median frequencies of these five genotypes is labelled the median simulation.

**Supplementary Table 1.** Summary of cases when MFT generates more MDR risk for the ASAQ double resistant genotype (*pfkelch13* 580Y, *pfcrt* 76T, *pfmdr1* 86Y Y184).

| $PfPR_{2-10}$ | Treatment Coverage | Importation | Median number of MDR risk-days (AUC) | Median AUC from adaptive MFT approach (50/50 AL and DHA-PPQ deployment) |
|---|---|---|---|---|
| 0.1% | 20% | None | MFT: 4.30<br>5yr Cyc: 3.85<br>Ad Cyc: 2.28 | Ad MFT: 1.22 |
| 0.1% | 40% | None | MFT: 4.93<br>5yr Cyc: 3.70<br>Ad Cyc: 3.56 | Ad MFT: 0.88 |
| 0.1% | 60% | None | MFT: 0.23<br>5yr Cyc: 0.11<br>Ad Cyc: 0.07 | Ad MFT: 0.06<br>( note that in this scenario malaria is eliminated after ten years ) |
| 1% | 20% | None | MFT: 5.33<br>Ad Cyc: 3.84 | Ad MFT: 2.95 |
| 1% | 60% | None | MFT: 2.19<br>5yr Cyc: 2.15<br>Ad Cyc: 2.16 | Ad MFT: 0.79 |
| 0.1% | 20% | One MDR genotype per year | MFT: 6.74<br>5yr Cyc: 6.43<br>Ad Cyc: 5.15 | Ad MFT: 2.11 |
| 0.1% | 60% | One MDR genotype per year | MFT: 212.22<br>Ad Cyc: 182.79 | MFT: 206.29<br>( note that this is a near-extinction scenarios with tens of infections present, and importation playing an outsized role in determining genotype frequencies ) |
| 1% | 20% | One MDR genotype per year | MFT: 7.00<br>5yr Cyc: 6.44<br>Ad Cyc: 4.71 | Ad MFT: 3.67<br>( but MDR risk of DHA-PPQ double-resistant becomes worse ) |

**Supplementary Table 2.** Number of mutations to maximally-resistant MDR types defined in Table 1 of main text. The mutation counts below are from the 'median simulation' (see Section 2 for definition) of a $PfPR_{2\text{-}10}$ = 5% setting with 40% treatment coverage and no importation. In the 5-year cycling strategy, DHA-PPQ is used first, ASAQ second, AL third, and DHA-PPQ is deployed again in years 16-20.

| | | Years 0-5 | Years 6-10 | Years 11-15 | Years 16-20 |
|---|---|---|---|---|---|
| DHA-PPQ, AQ triple-resistant | MFT | 16 | 47 | 126 | 286 |
| | 5-year cycling | 47 | 110 | 103 | 396 |
| DHA-PPQ, LUM triple-resistant | MFT | 0 | 0 | 0 | 5 |
| | 5-year cycling | 0 | 0 | 0 | 0 |
| DHA-PPQ double-resistant | MFT | 29 | 128 | 314 | 789 |
| | 5-year cycling | 132 | 237 | 263 | 928 |
| ASAQ double-resistant | MFT | 790 | 730 | 849 | 947 |
| | 5-year cycling | 829 | 932 | 928 | 925 |
| AL double resistant | MFT | 0 | 0 | 0 | 6 |
| | 5-year cycling | 0 | 0 | 0 | 0 |

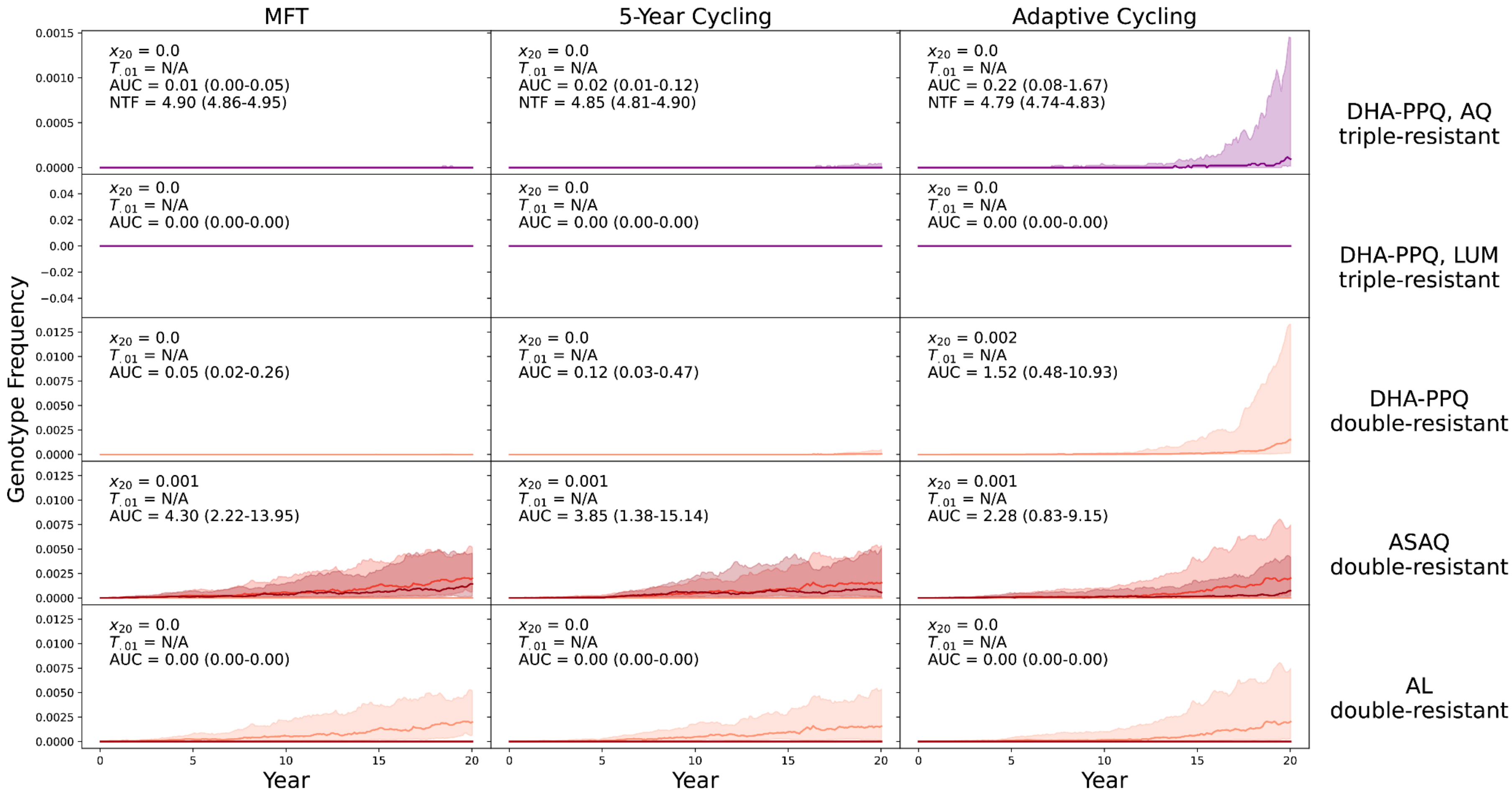


**Supplementary Figure 1.** PfPR$_{2\text{-}10}$ = 0.1% and treatment coverage = 20%. No importation. Each row shows the median trajectories, with shaded areas showing interquartile ranges, of the double-resistant or triple-resistant labelled at the right. In the bottom two rows, dark red corresponds to quadruple-mutant double-resistance, medium red corresponds to triple-mutant double-resistance, and light red corresponds to double-mutant double-resistance. The columns correspond to the three different drug deployment strategies. The final mutant genotype frequency after 20 years ($x_{20}$), the time until this genotype reaches 0.01 frequency ($T_{.01}$), the number of MDR risk-days for this mutant (AUC), and the number of treatment failures (NTF) per 100 persons per year are shown in each panel. Note that for the ASAQ double-resistant, MFT has a higher AUC value than either cycling strategy.

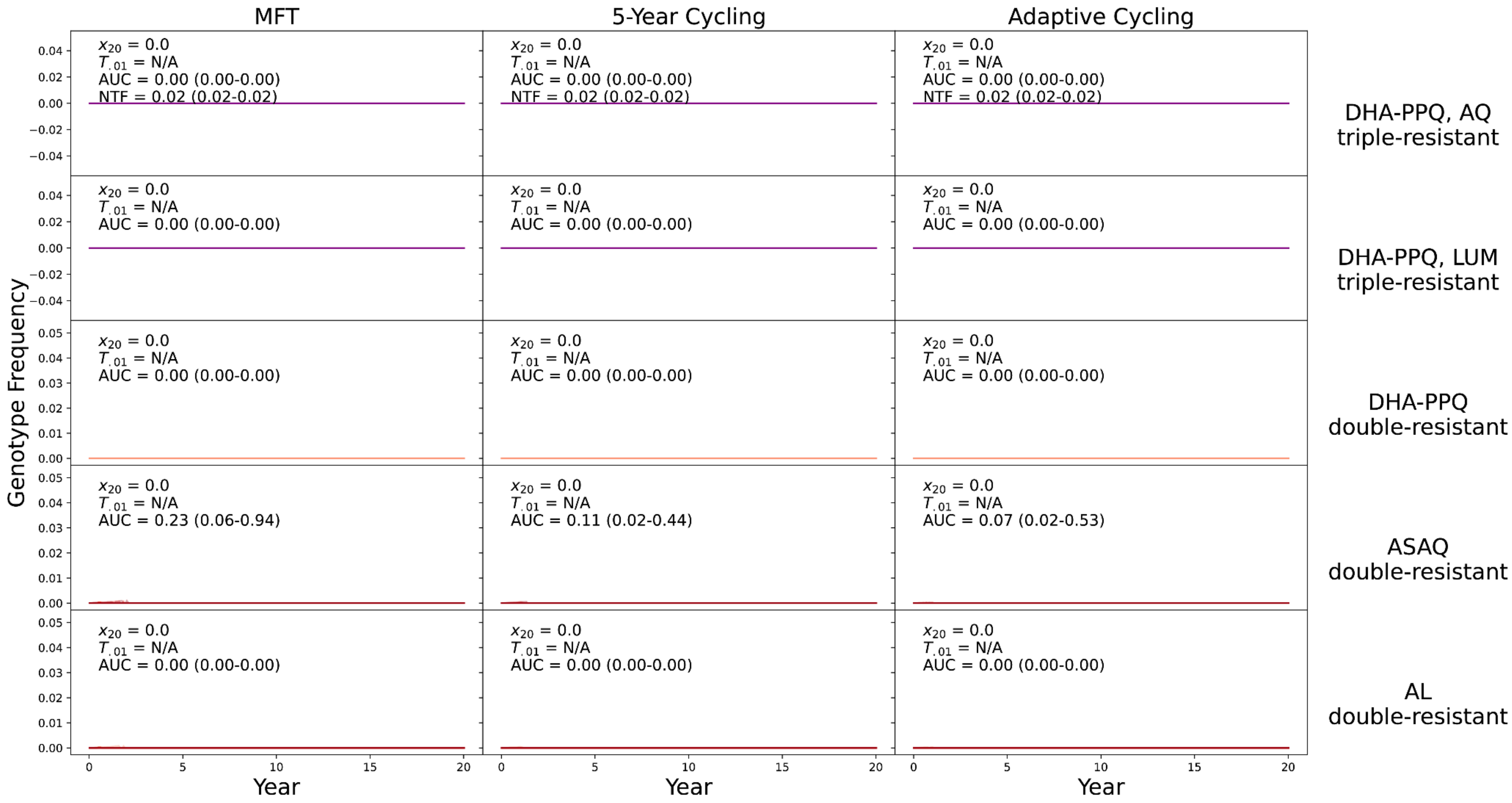


**Supplementary Figure 2.** PfPR$_{2-10}$ = 0.1% and treatment coverage = 60%. No importation. 40% coverage is shown in Figure 3 of the main text. Note that for the ASAQ double-resistant, MFT has a higher AUC value than either cycling strategy. In these runs, malaria is effectively eliminated in more than 95% of simulations, for all strategies. The AUC values show the sums, in units of frequency-days, during years 0 to 10, prior to elimination.

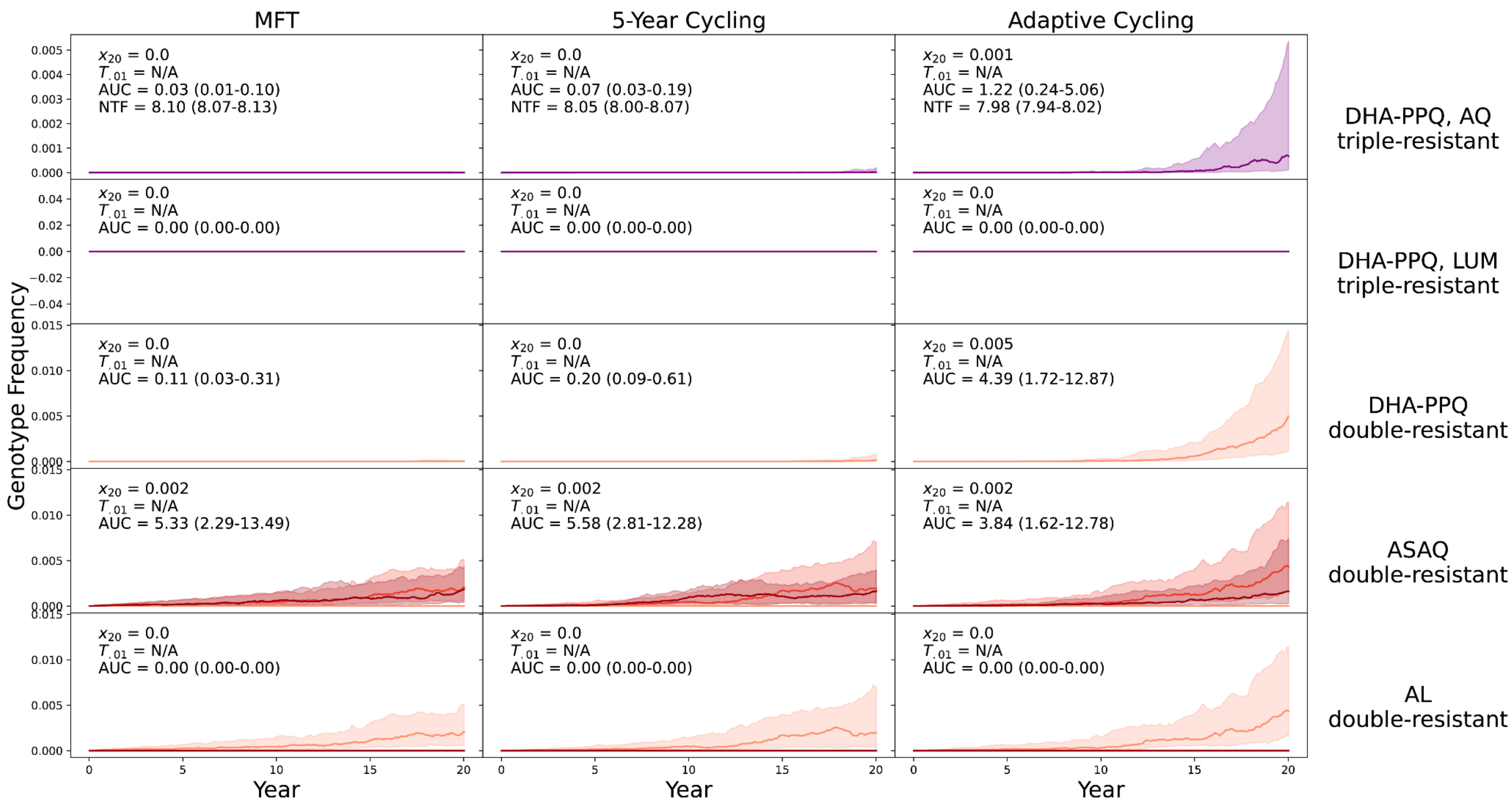


**Supplementary Figure 3.** PfPR$_{2\text{-}10}$ = 1% and treatment coverage = 20%. No importation. Note that for the ASAQ double-resistant, MFT has a higher AUC value than the adaptive cycling strategy.

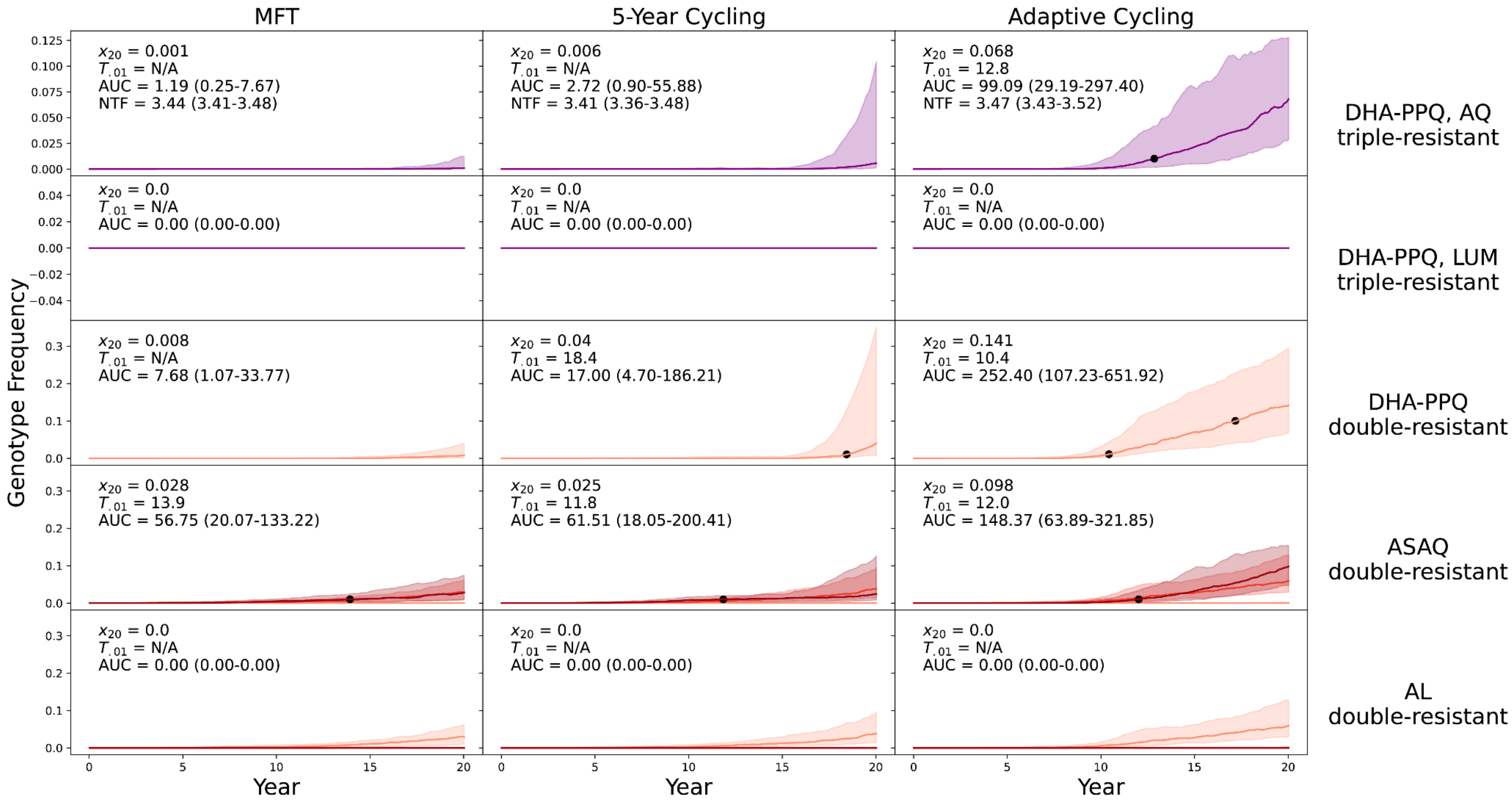


**Supplementary Figure 4.** $PfPR_{2-10}$ = 1% and treatment coverage = 40%. No importation.

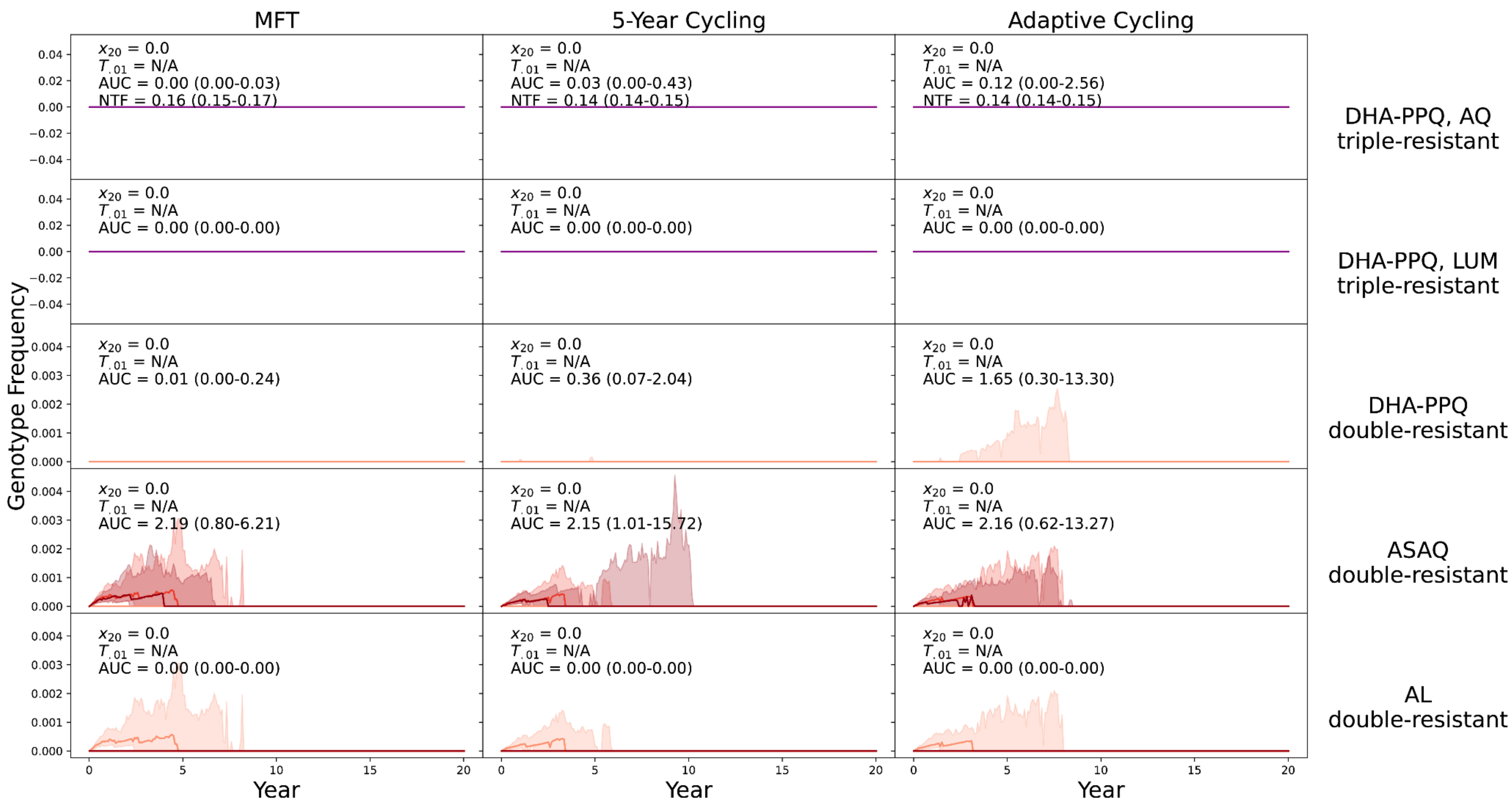


**Supplementary Figure 5.** PfPR$_{2\text{-}10}$ = 1% and treatment coverage = 60%. No importation. Note that for the ASAQ double-resistant, MFT has a higher AUC value than either cycling strategy.

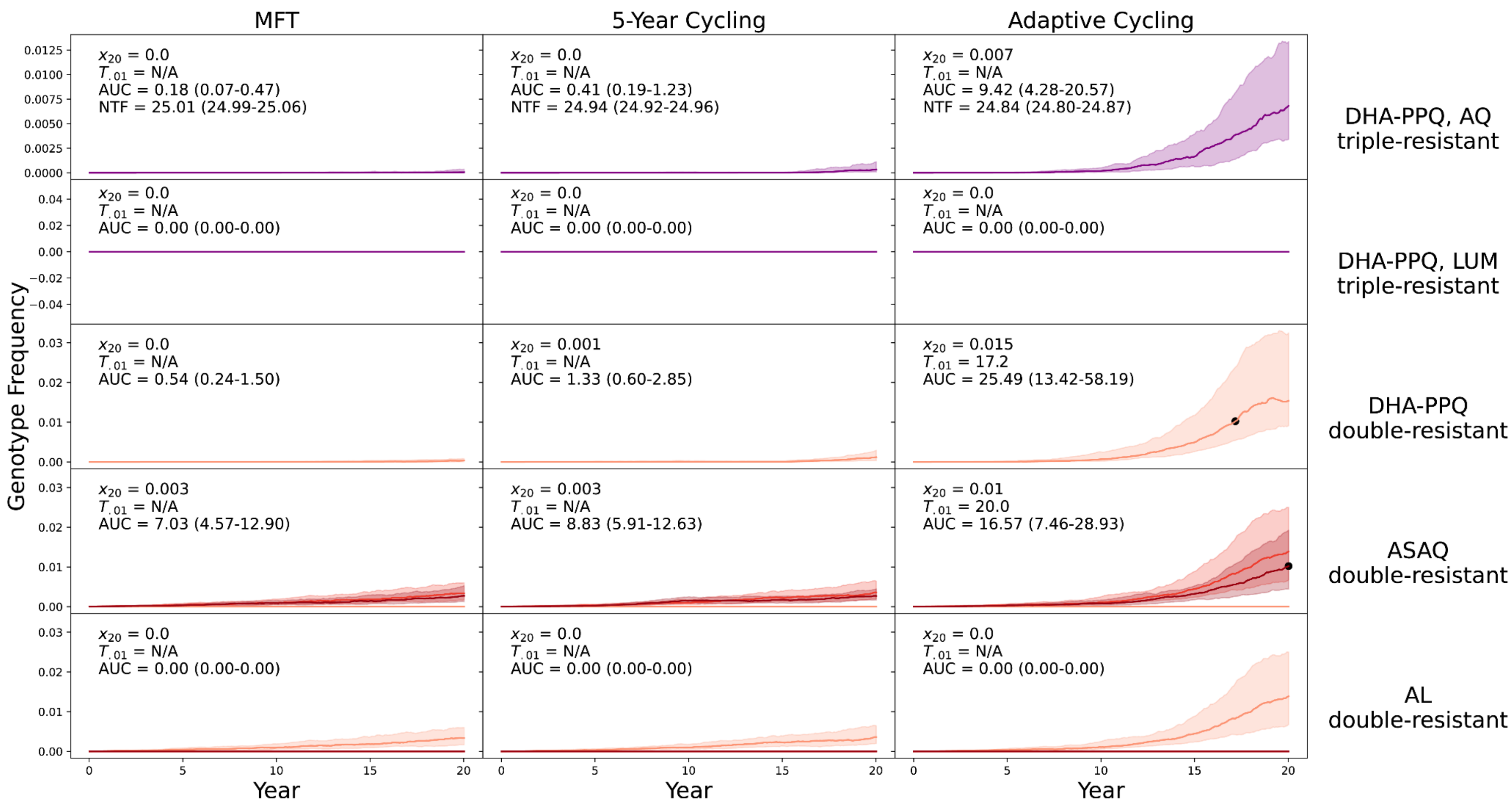


**Supplementary Figure 6.** PfPR$_{2\text{-}10}$ = 5% and treatment coverage = 20%. No importation.

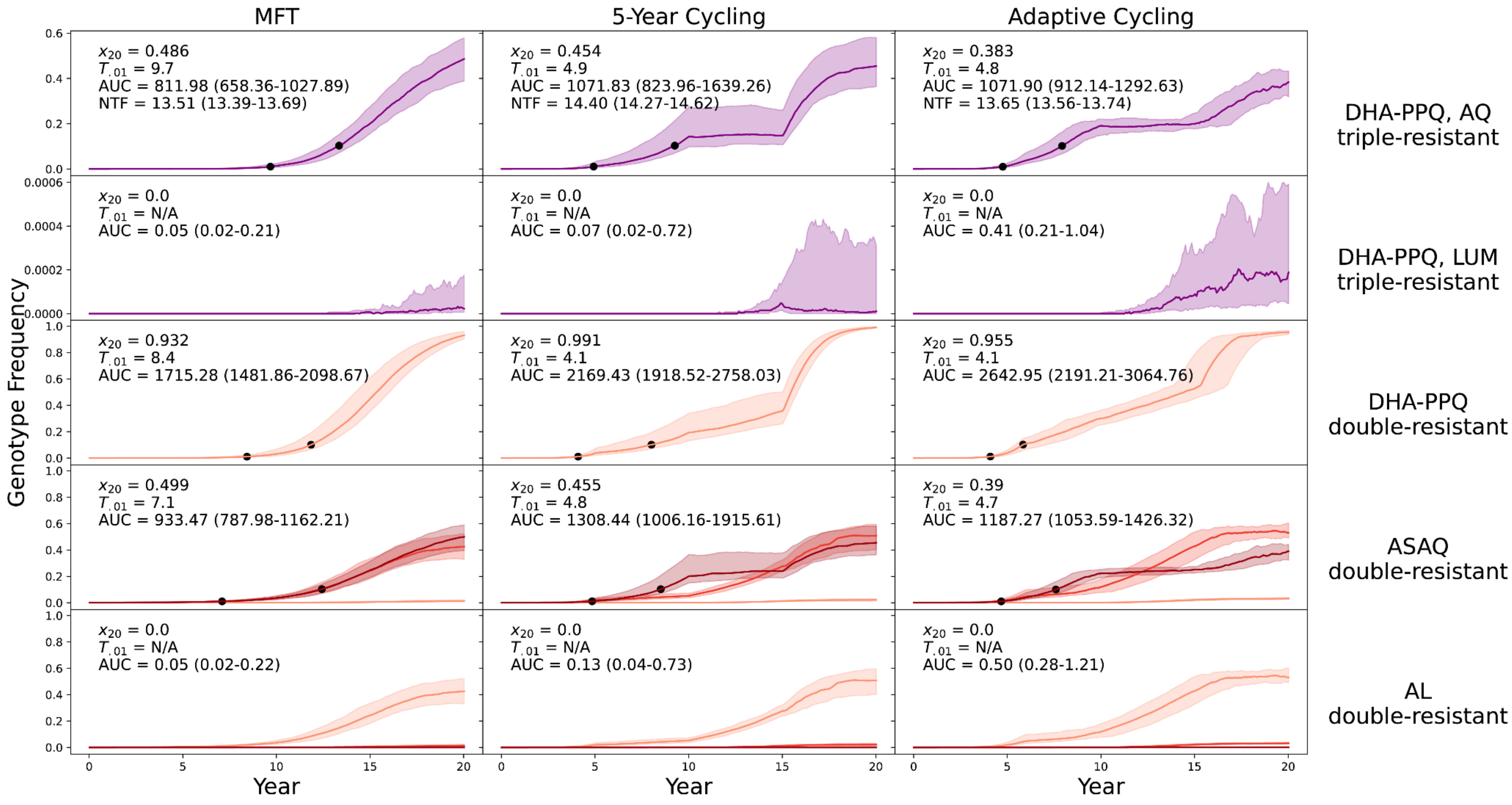


**Supplementary Figure 7.** PfPR$_{2\text{-}10}$ = 5% and treatment coverage = 60%. No importation. 40% coverage shown in Figure 2 of main text.

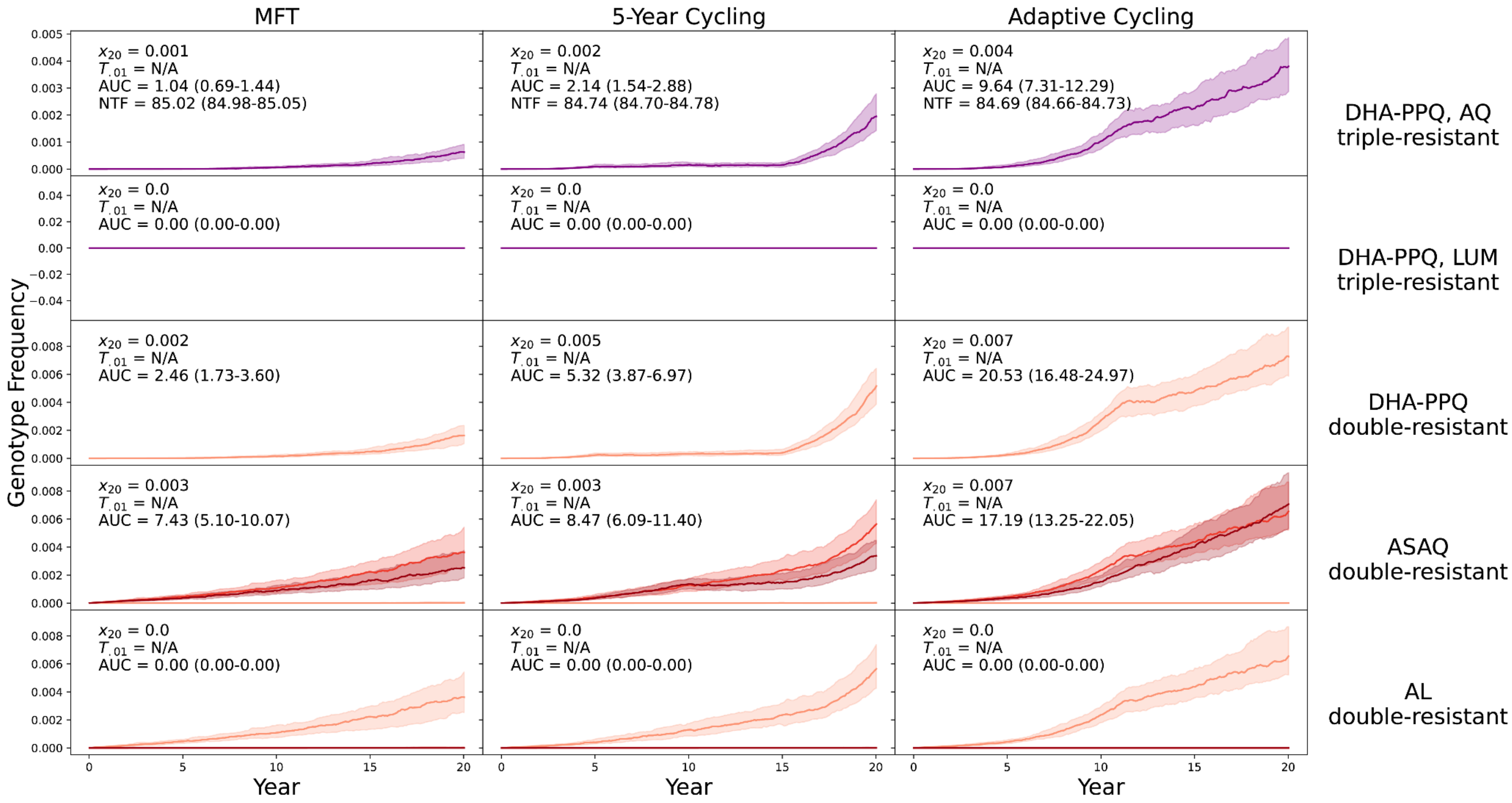


**Supplementary Figure 8.** PfPR$_{2\text{-}10}$ = 25% and treatment coverage = 20%. No importation.

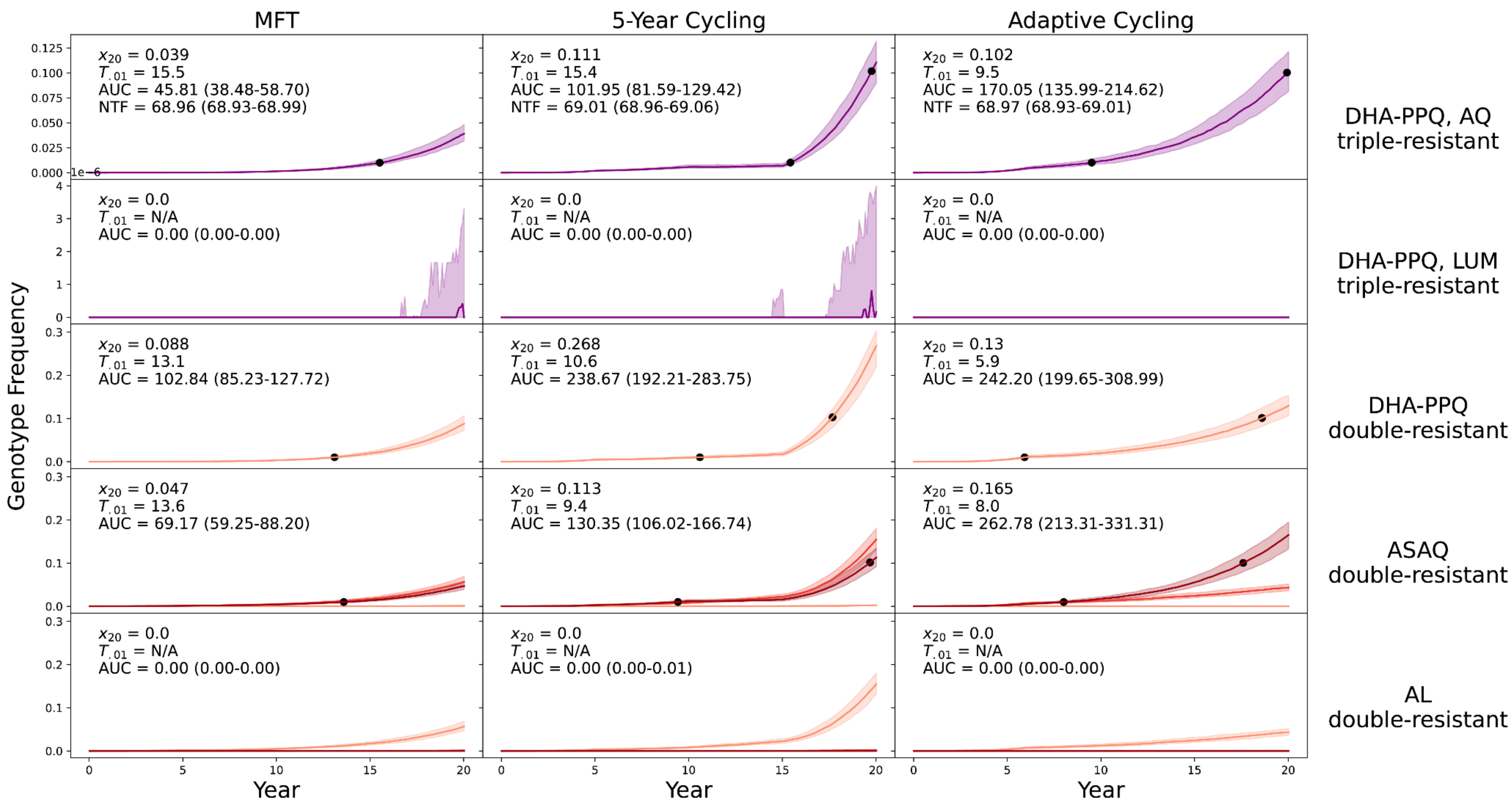


**Supplementary Figure 9.** PfPR$_{2\text{-}10}$ = 25% and treatment coverage = 40%. No importation.

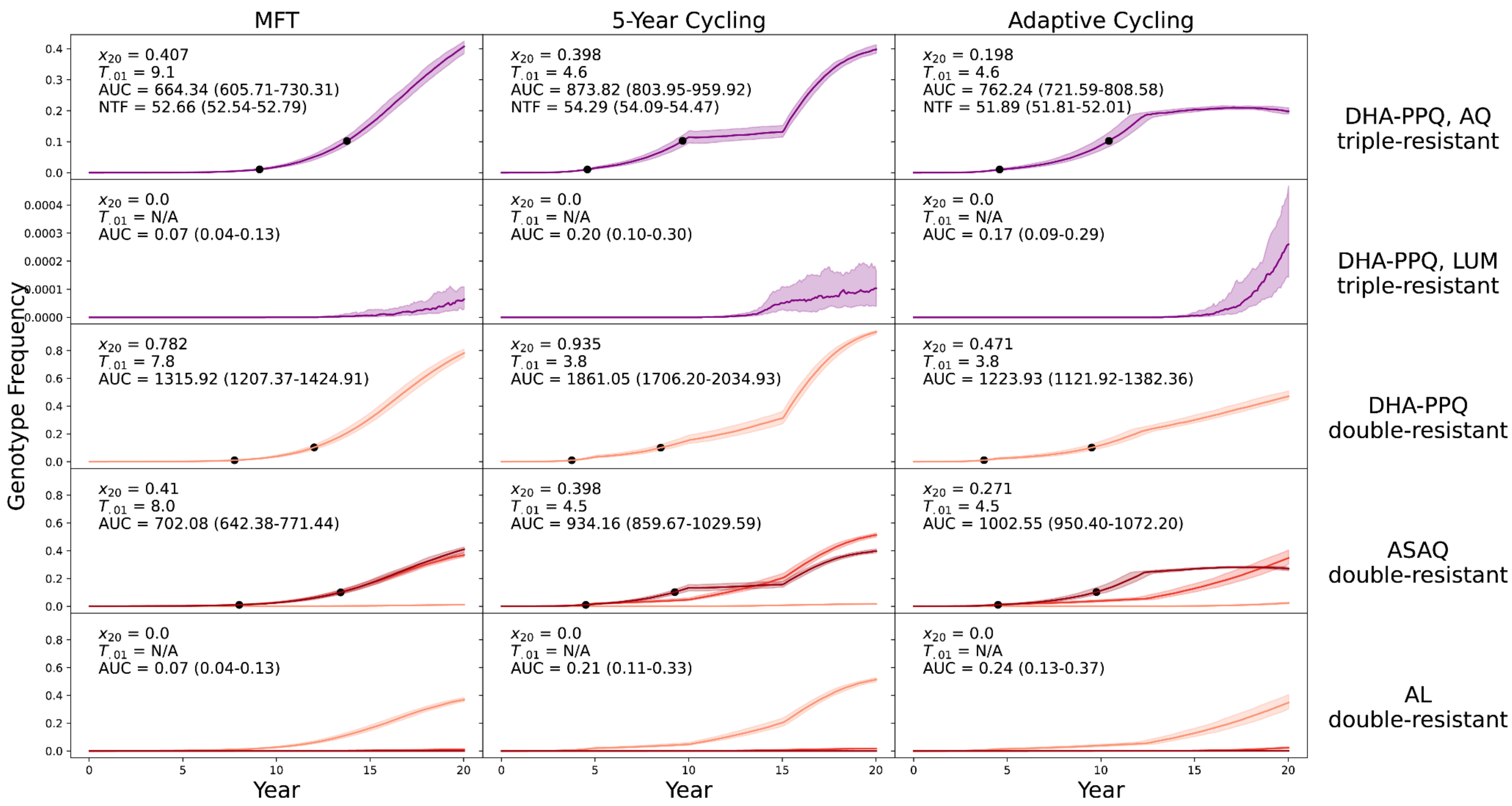


**Supplementary Figure 10.** PfPR$_{2\text{-}10}$ = 25% and treatment coverage = 60%. No importation. Note that for the DHA-PPQ double-resistant, MFT has a higher AUC value than the adaptive cycling strategy.

### Twelve epidemiological scenarios with importation (summarized in Figure 5 of main text)

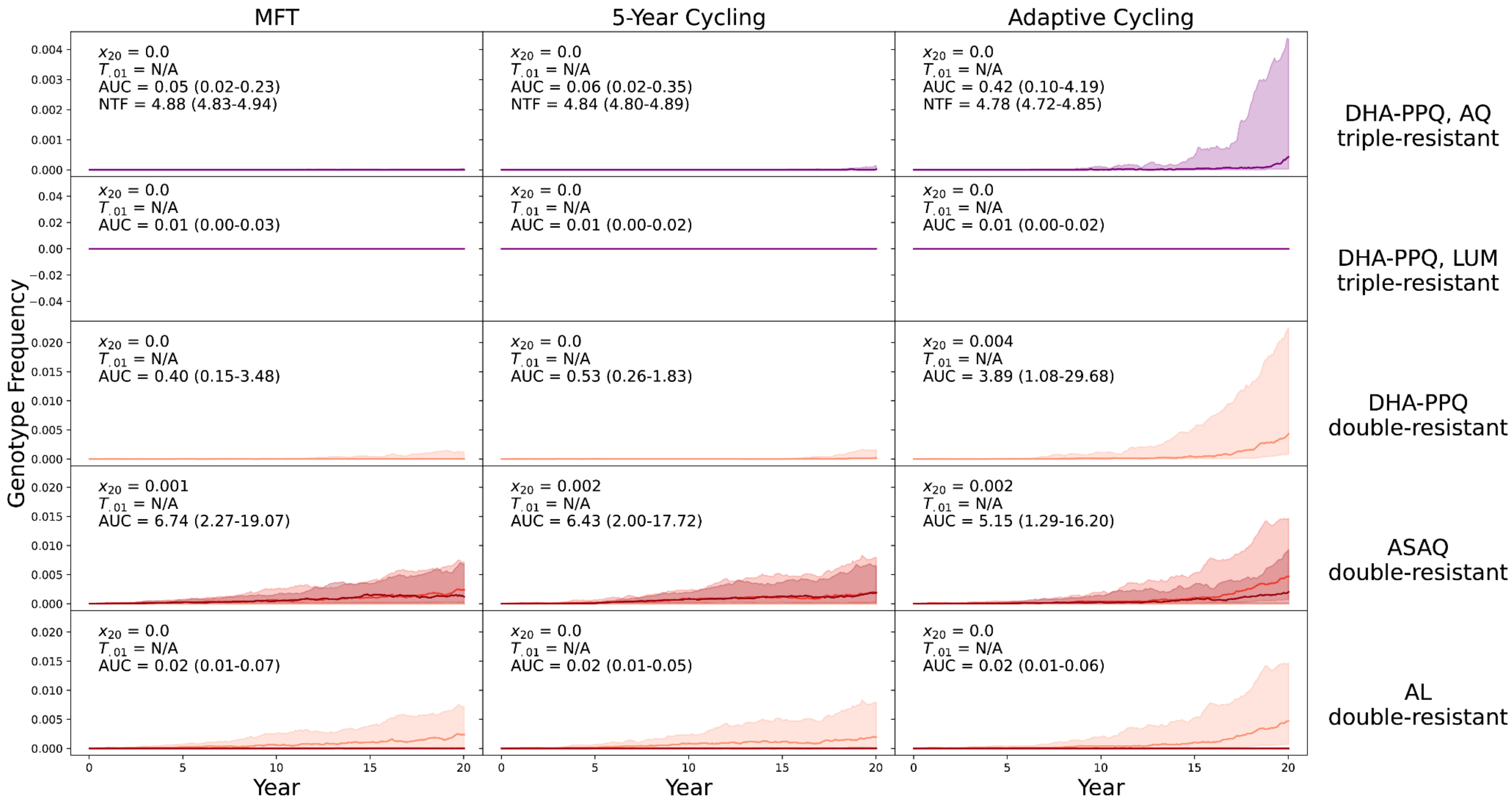


**Supplementary Figure 11.** $PfPR_{2\text{-}10}$ = 0.1% and treatment coverage = 20%. With Importation. Note that for the ASAQ double-resistant, MFT has a higher AUC value than either cycling strategy.

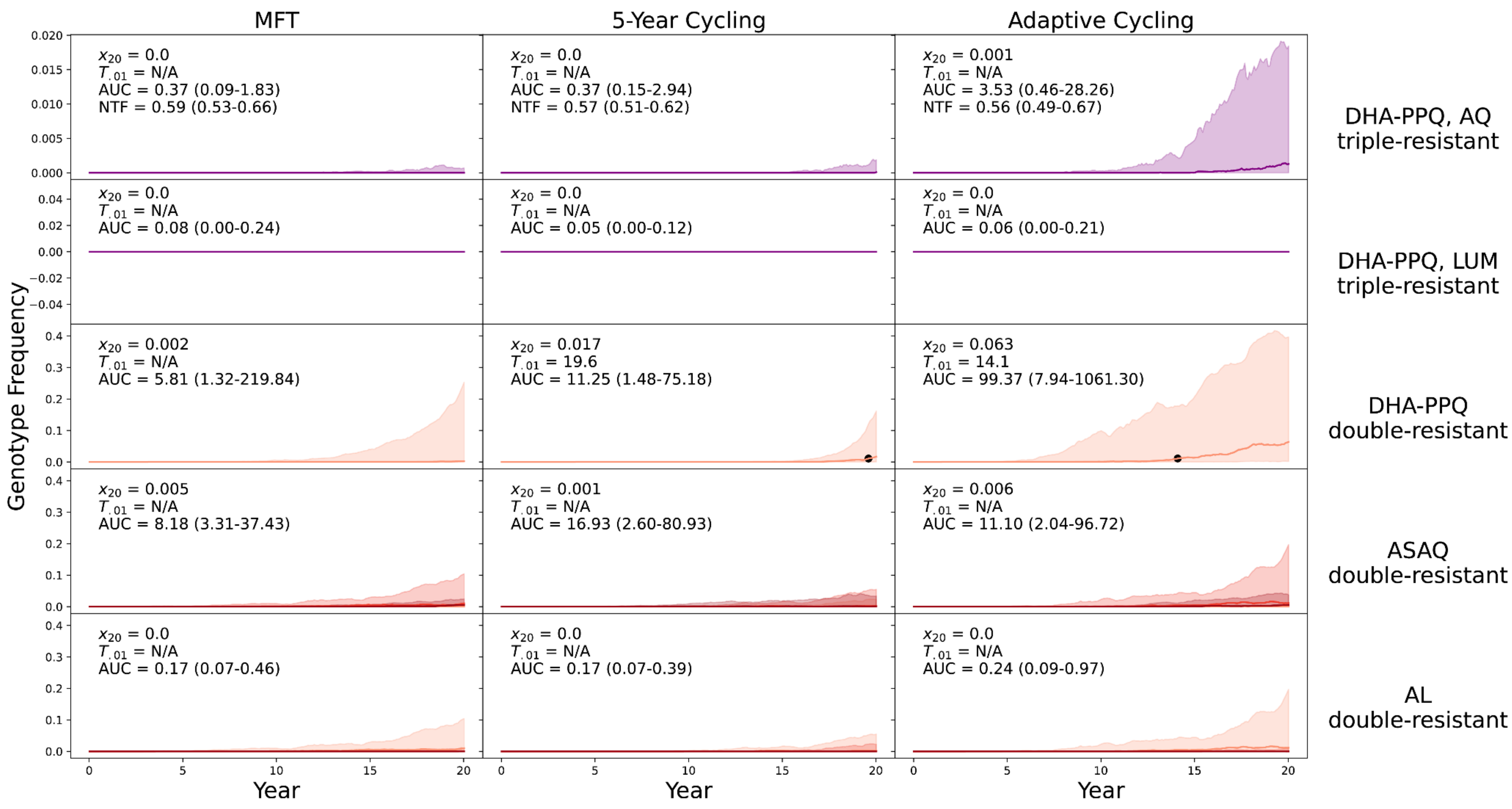


**Supplementary Figure 12.** PfPR$_{2\text{-}10}$ = 0.1% and treatment coverage = 40%. With Importation. Note that for the DHA-PPQ-LUM triple-resistant, MFT has a higher AUC value than either cycling strategy.

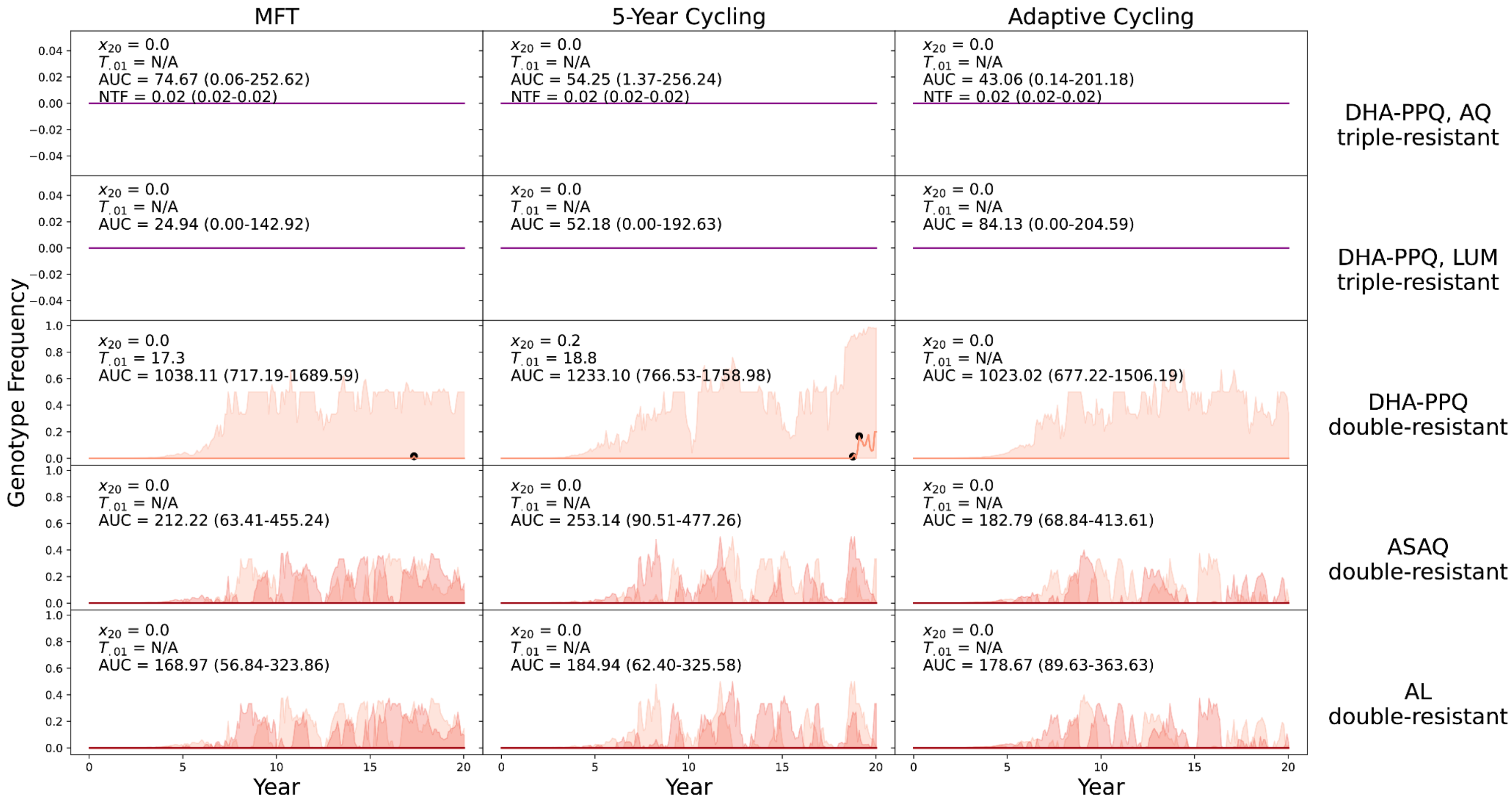


**Supplementary Figure 13.** PfPR$_{2\text{-}10}$ = 0.1% and treatment coverage = 60%. With Importation. Note that in 4 out of 10 comparisons MFT has a higher AUC value than cycling. This high-coverage low-prevalence scenario has the majority of its simulations runs reach extinction levels (PfPR$_{2\text{-}10}$ < 0.01% for more than 95% of runs) which corresponds to double-digit counts of parasite positive individuals. For this reason, the MDR frequencies are sometimes very high because they are being imported into a low case number environment, and the AUC values are much higher than in Supplementary Figure 12 or Supplementary Figure 16.

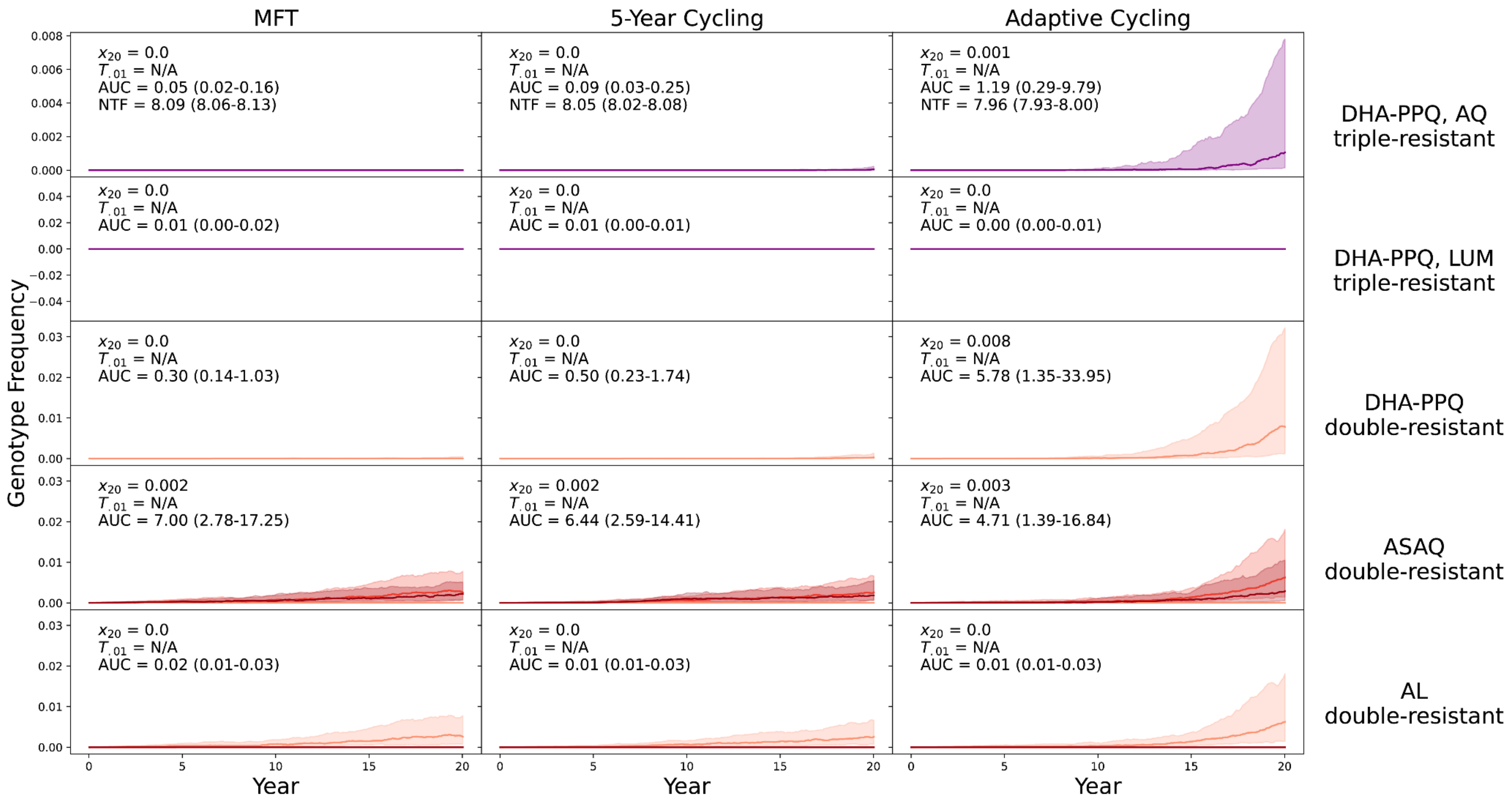


**Supplementary Figure 14.** $PfPR_{2-10}$ = 1% and treatment coverage = 20%. With Importation. Note that in 5 out of 10 comparisons MFT has a higher AUC value than cycling.

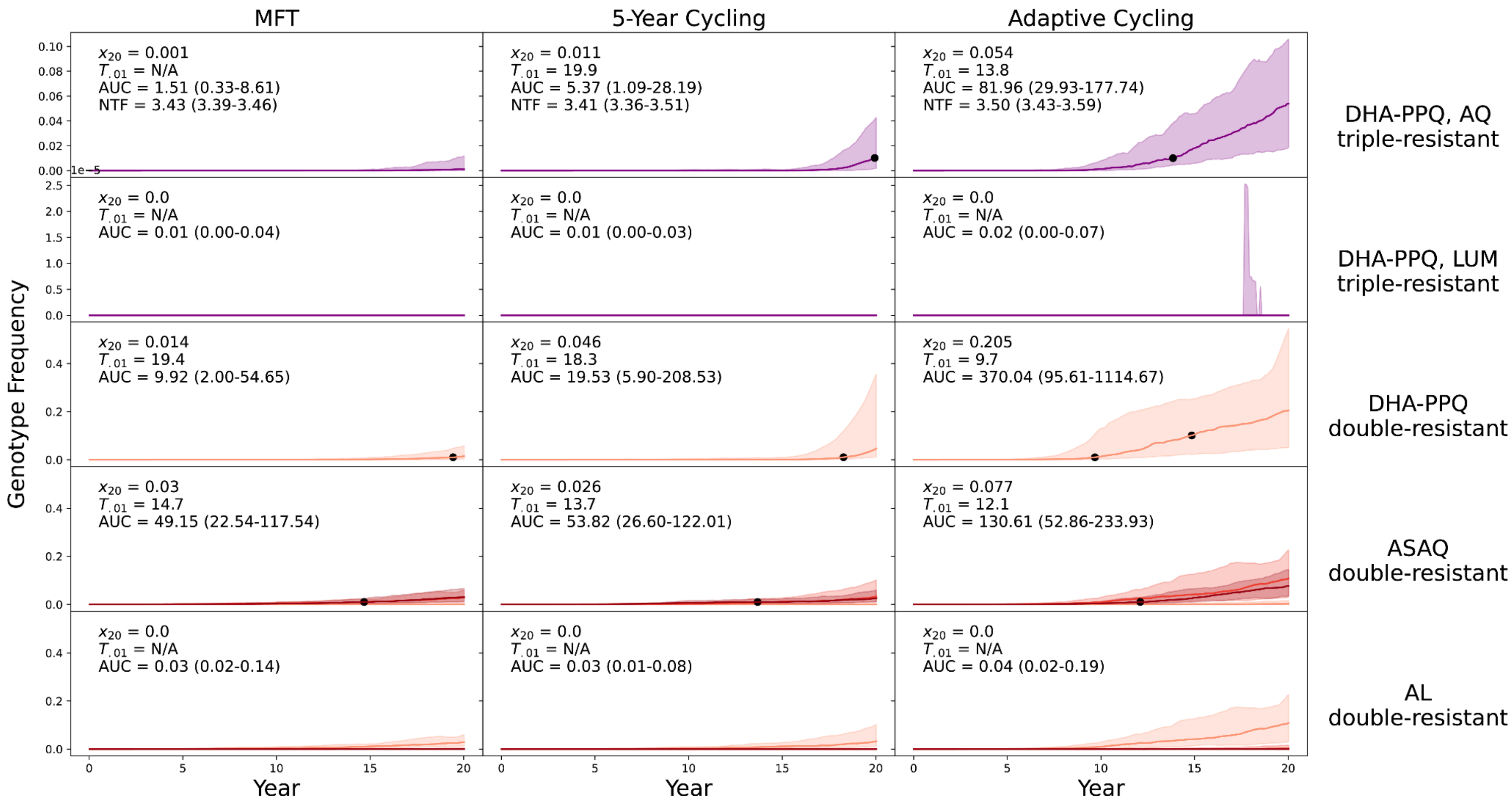


**Supplementary Figure 15.** $PfPR_{2\text{-}10}$ = 1% and treatment coverage = 40%. With Importation.

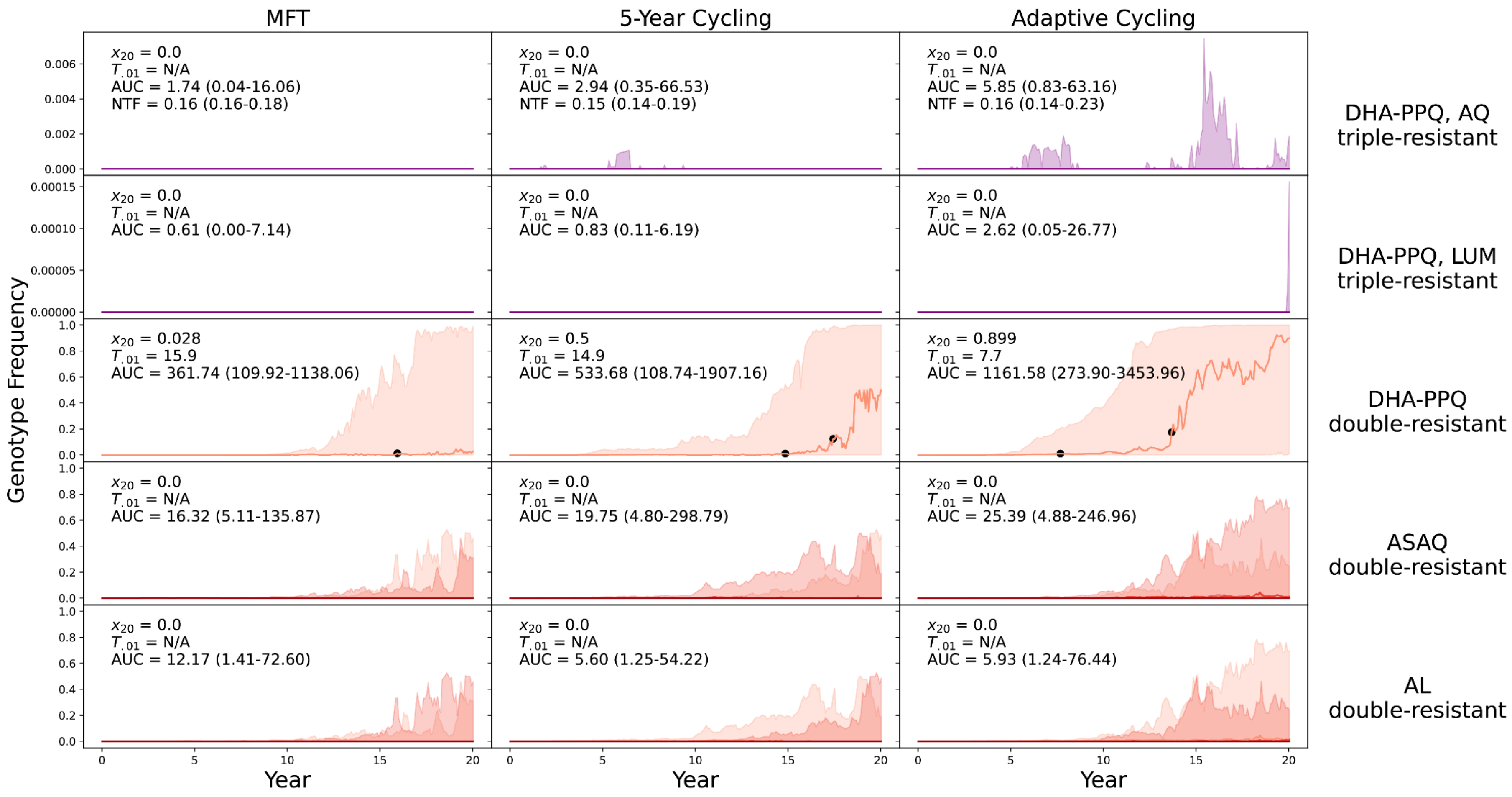


**Supplementary Figure 16.** PfPR$_{2\text{-}10}$ = 1% and treatment coverage = 60%. With Importation. Note that for the AL double-resistant, MFT has a higher AUC value than either cycling strategy.

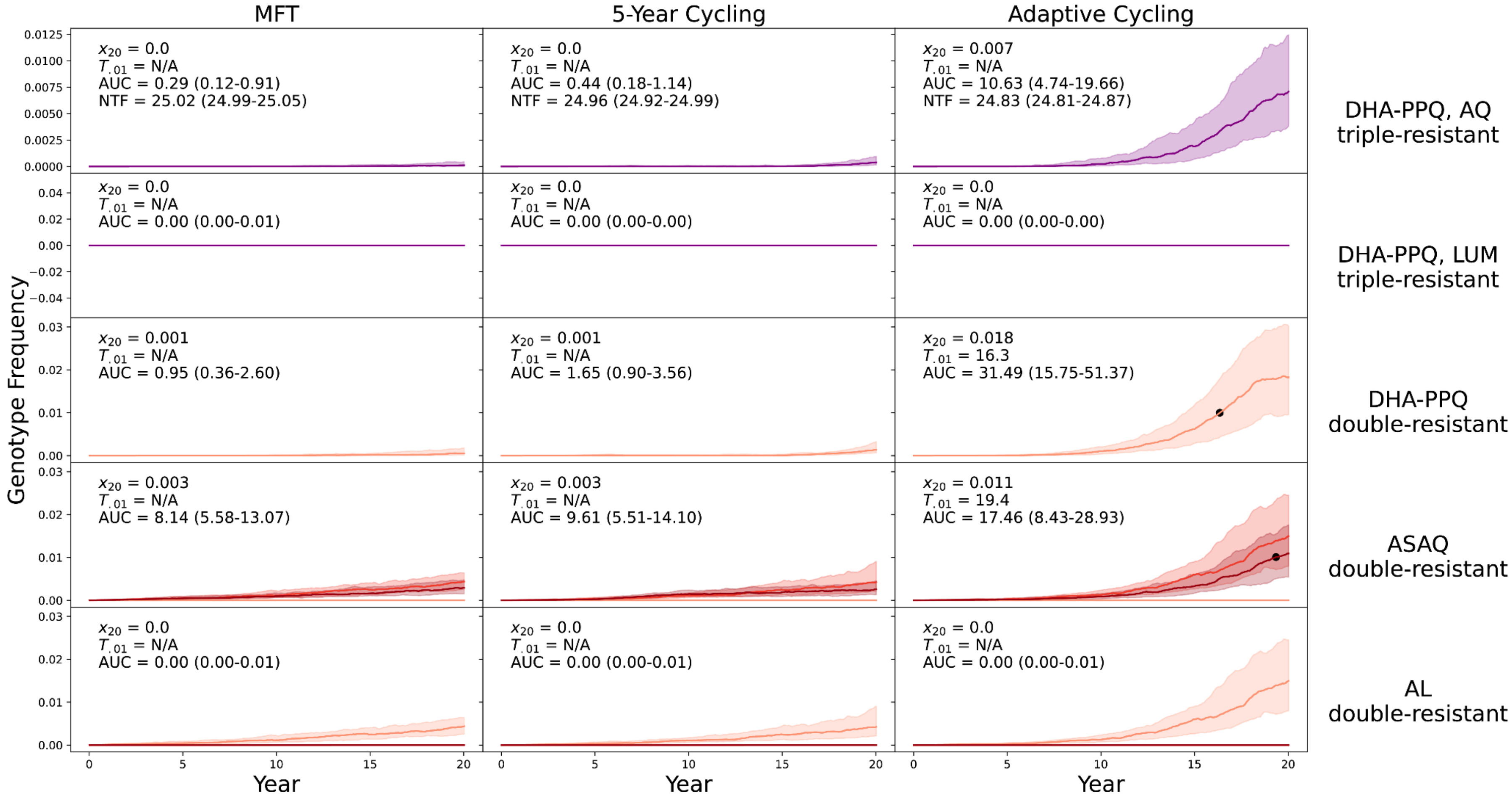


**Supplementary Figure 17.** PfPR$_{2\text{-}10}$ = 5% and treatment coverage = 20%. With Importation.

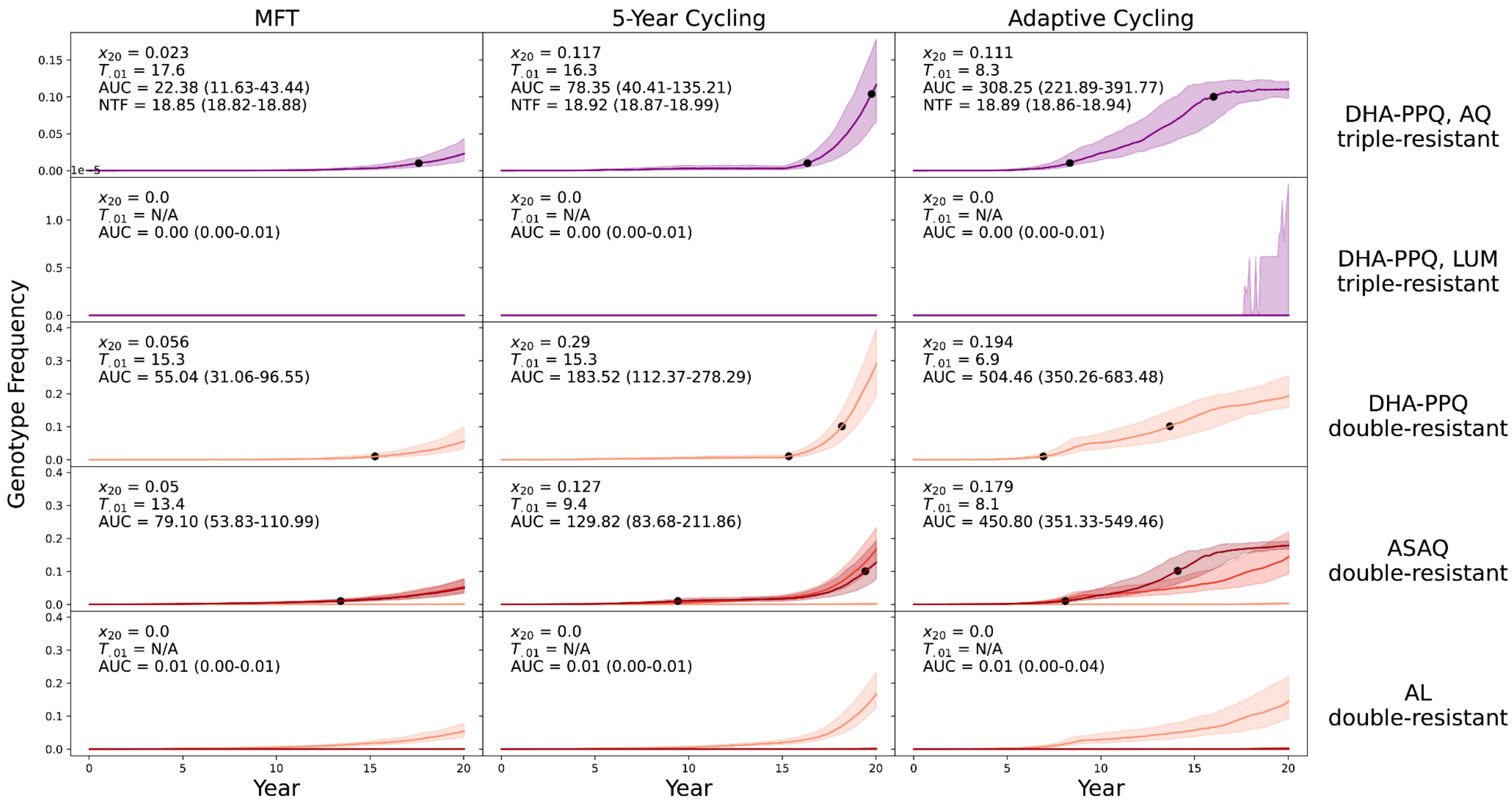


**Supplementary Figure 18.** PfPR$_{2\text{-}10}$ = 5% and treatment coverage = 40%. With Importation.

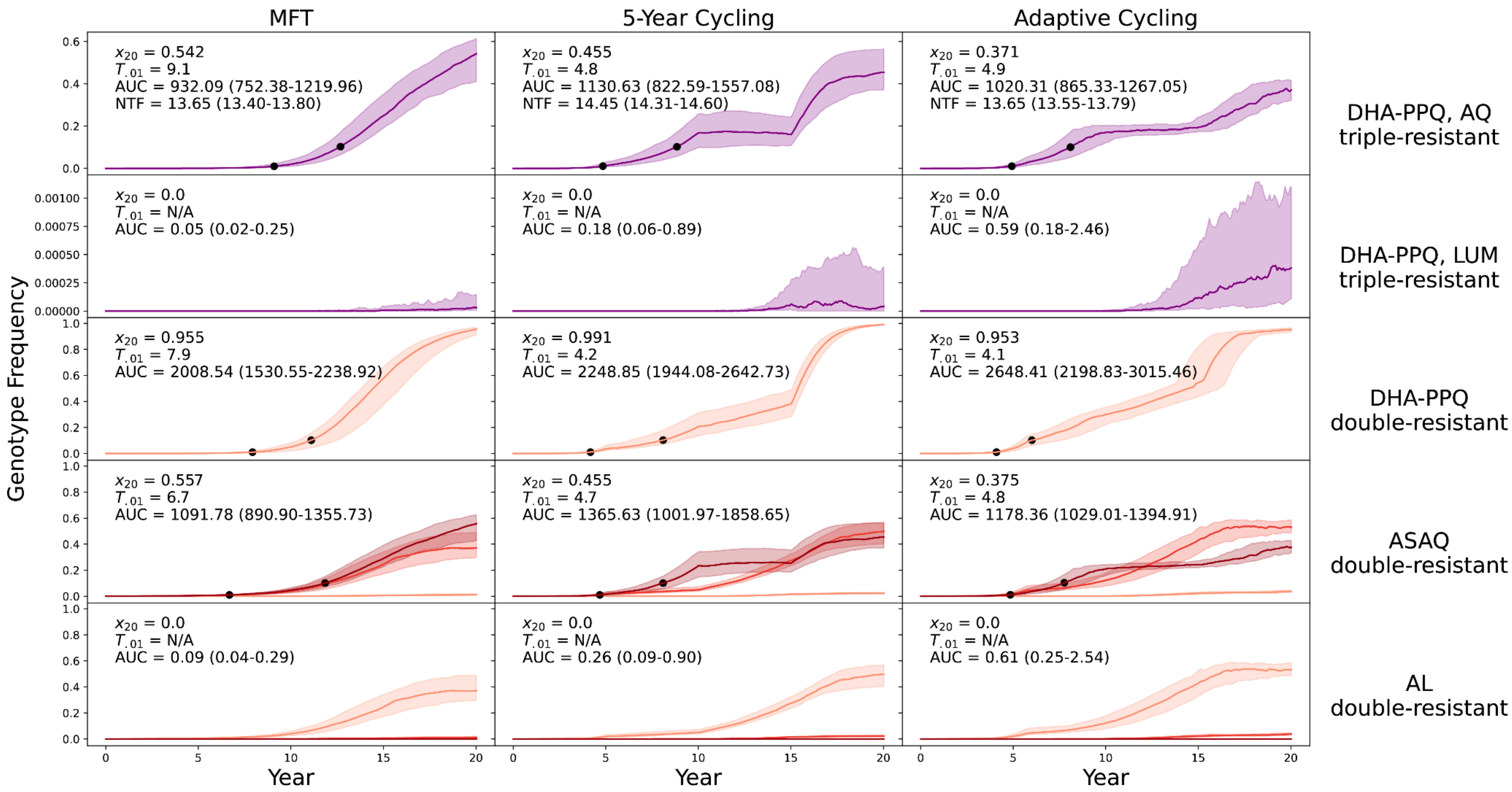


**Supplementary Figure 19.** PfPR$_{2-10}$ = 5% and treatment coverage = 60%. With Importation.

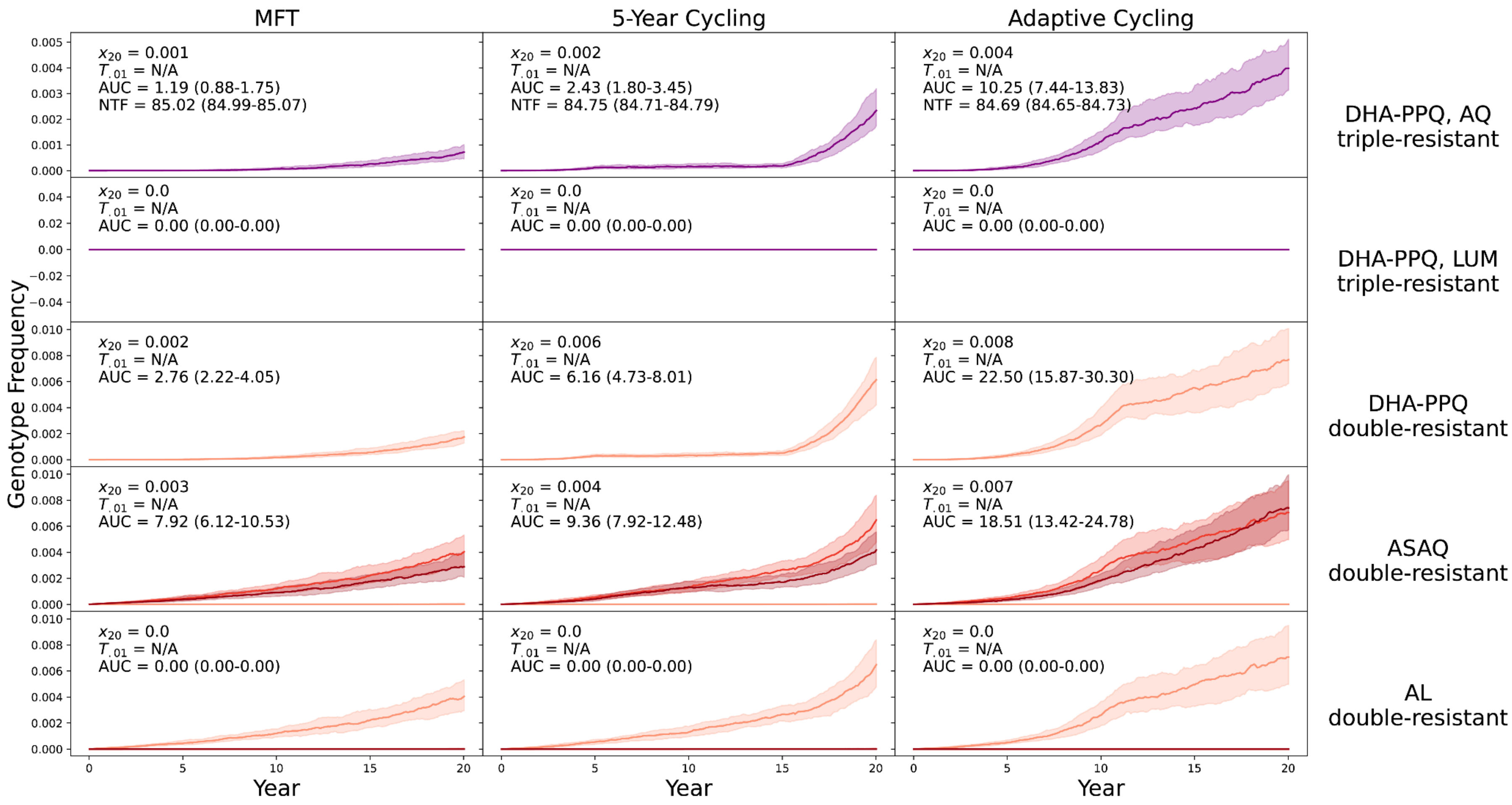


**Supplementary Figure 20.** PfPR$_{2-10}$ = 25% and treatment coverage = 20%. With Importation.

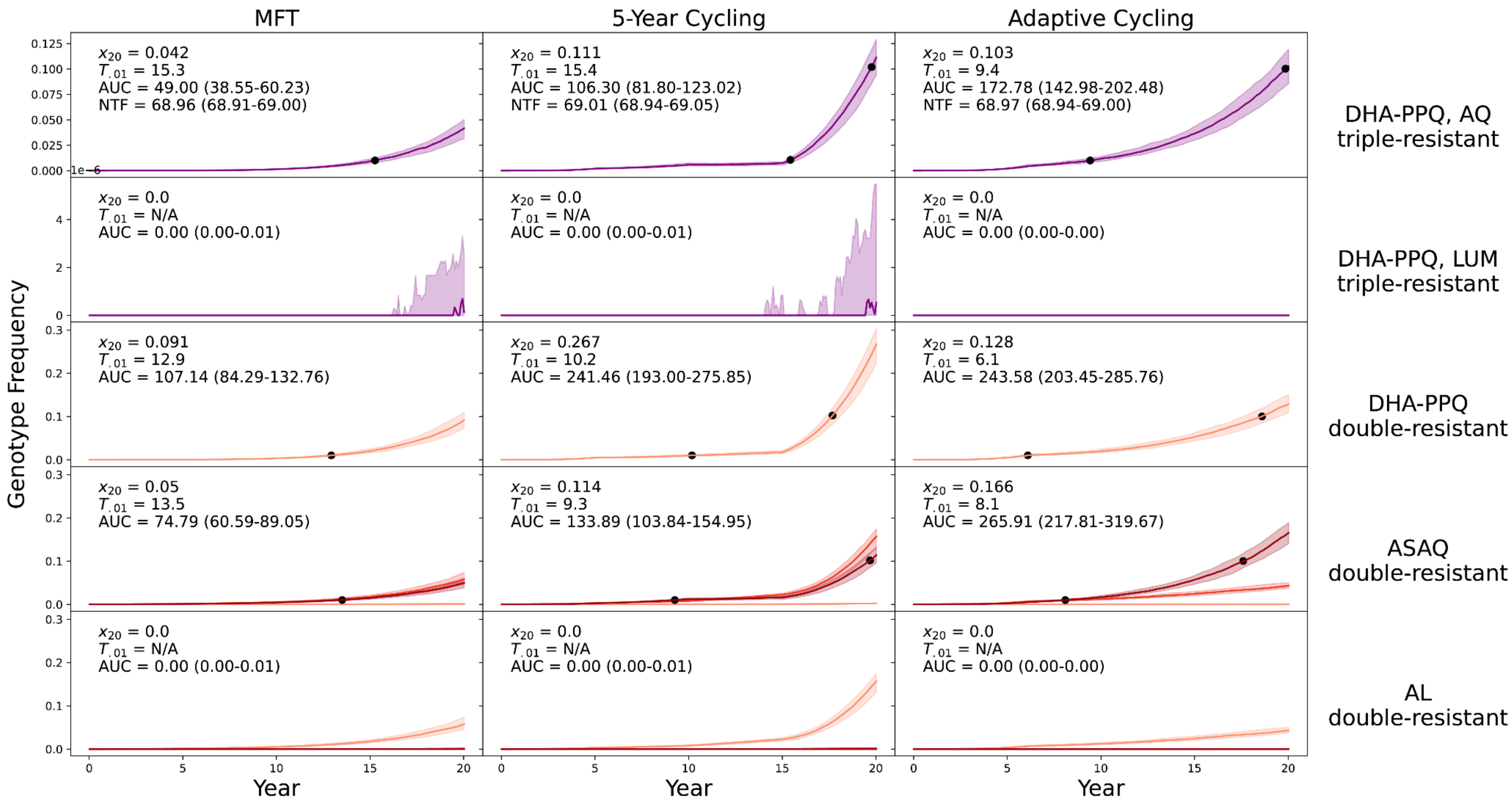


**Supplementary Figure 21.** PfPR$_{2\text{-}10}$ = 25% and treatment coverage = 40%. With Importation.

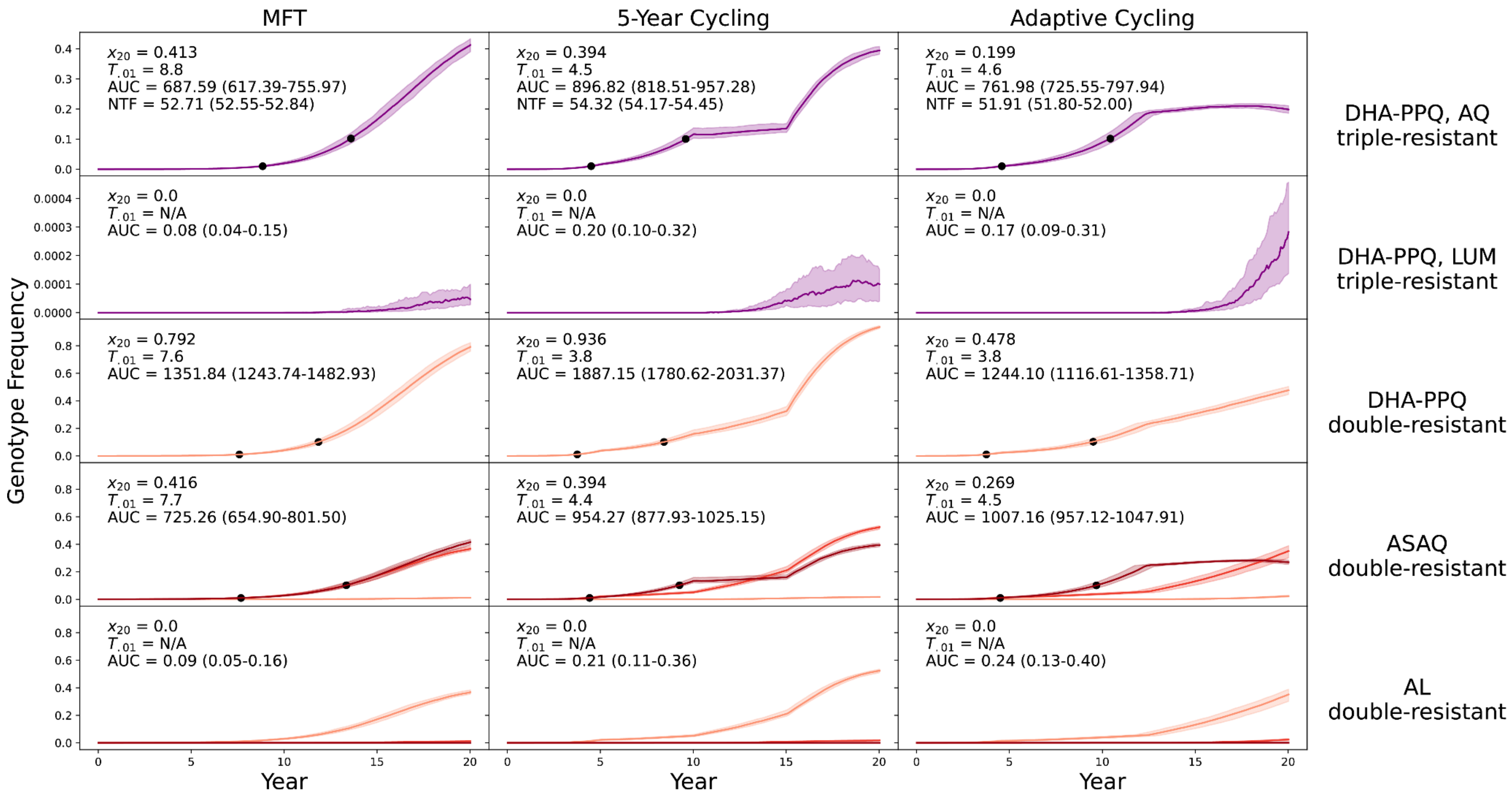


**Supplementary Figure 22.** PfPR$_{2\text{-}10}$ = 25% and treatment coverage = 60%. With Importation. Note that for the DHA-PPQ double-resistant, MFT has a higher AUC value than adaptive cycling.

### Two scenarios with lower mutation rate

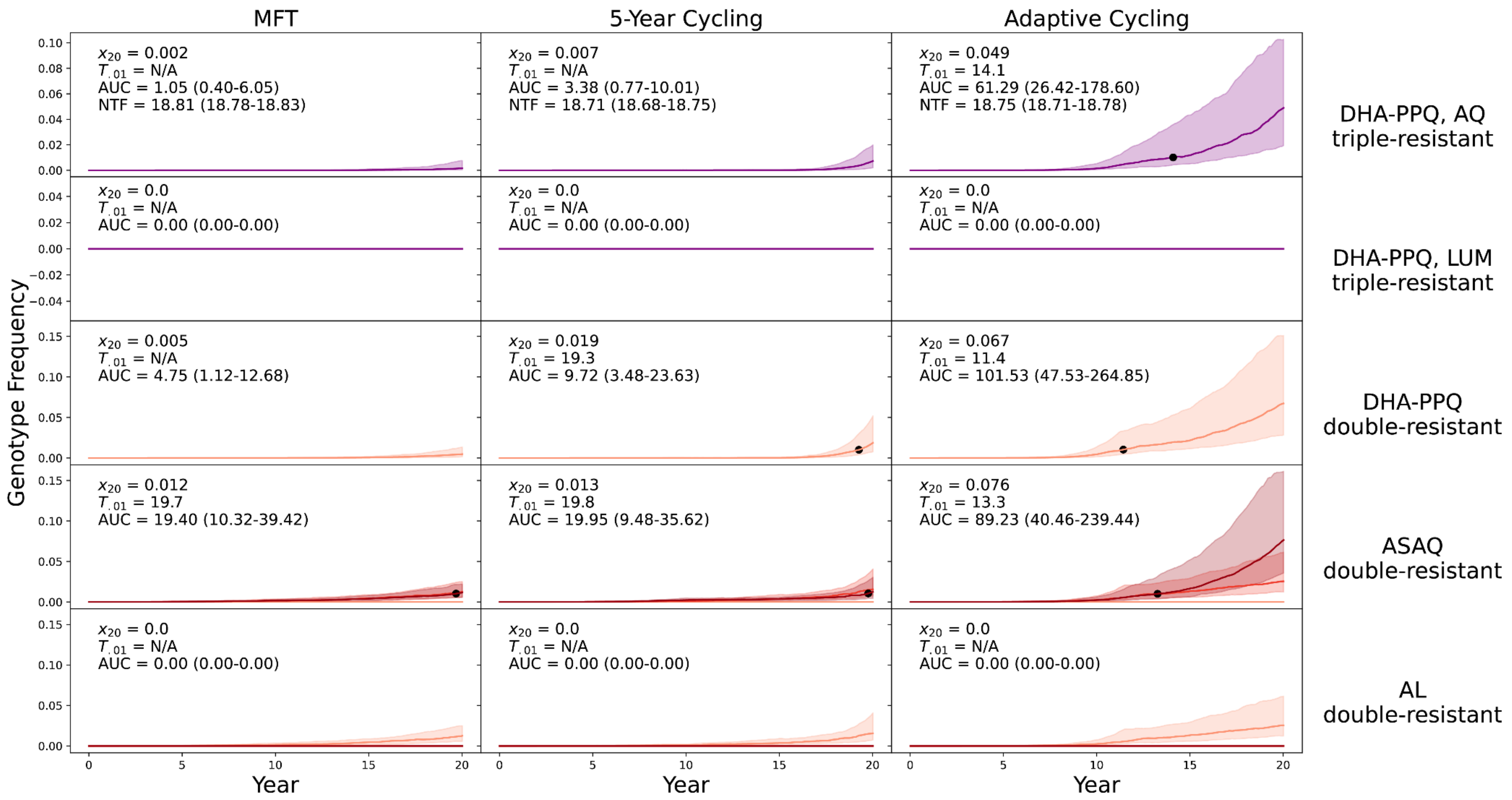


**Supplementary Figure 23.** PfPR$_{2\text{-}10}$ = 5% and treatment coverage = 40%. No importation. Mutation rate reduced 3-fold from 0.001983 per treated case to 0.000661 per treated case. Relationships among the strategies are similar but delay to resistance emergence is now >19 years for MFT and 5-year cycling.

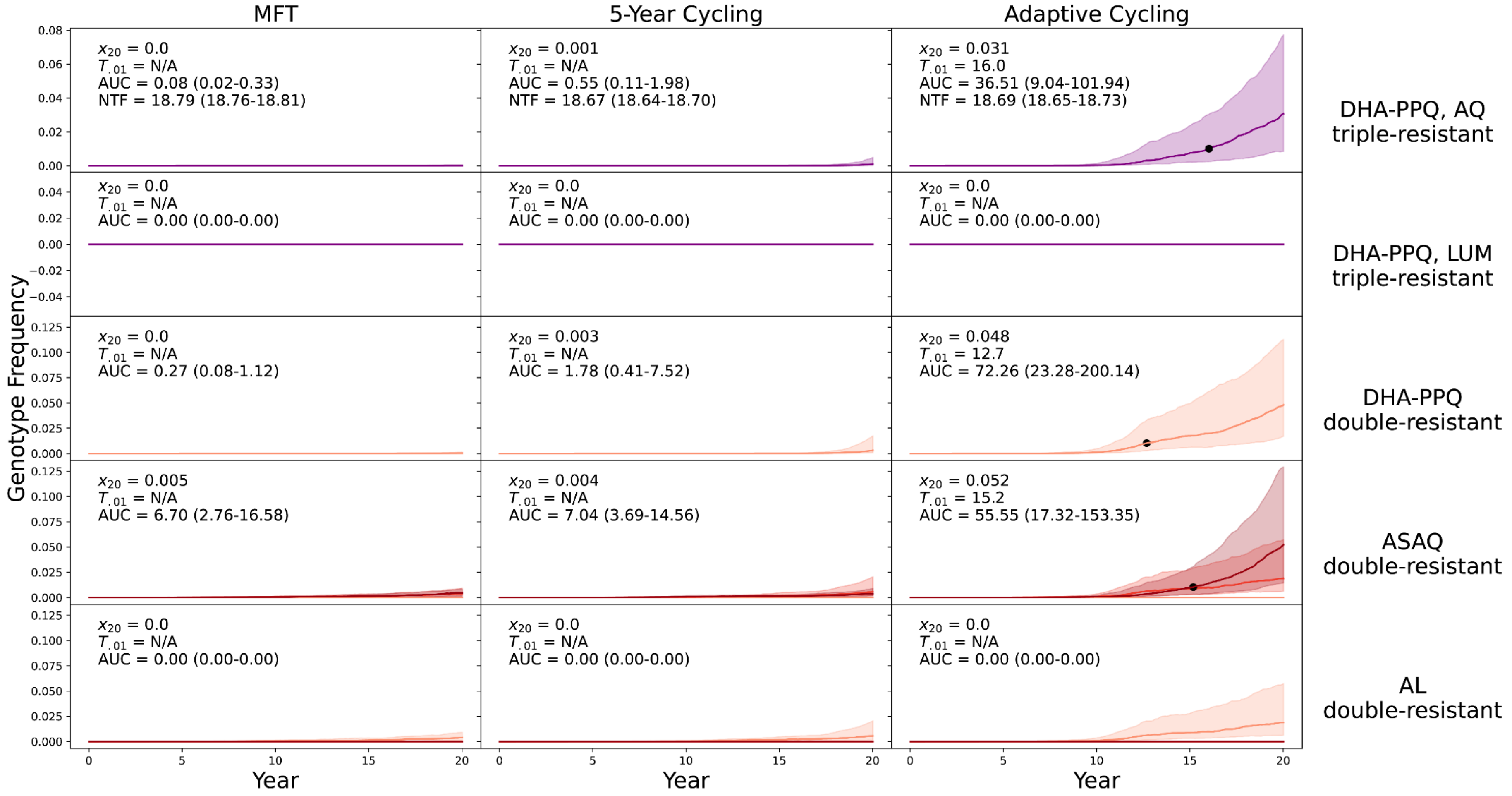


**Supplementary Figure 24.** PfPR$_{2\text{-}10}$ = 5% and treatment coverage = 40%. No importation. Mutation rate reduced 5-fold from 0.001983 per treated case to 0.0003966 per treated case. Relationships among the strategies are similar but delay to resistance emergence is now >20 years for MFT and 5-year cycling.

### Prevalence levels for all 12 scenarios

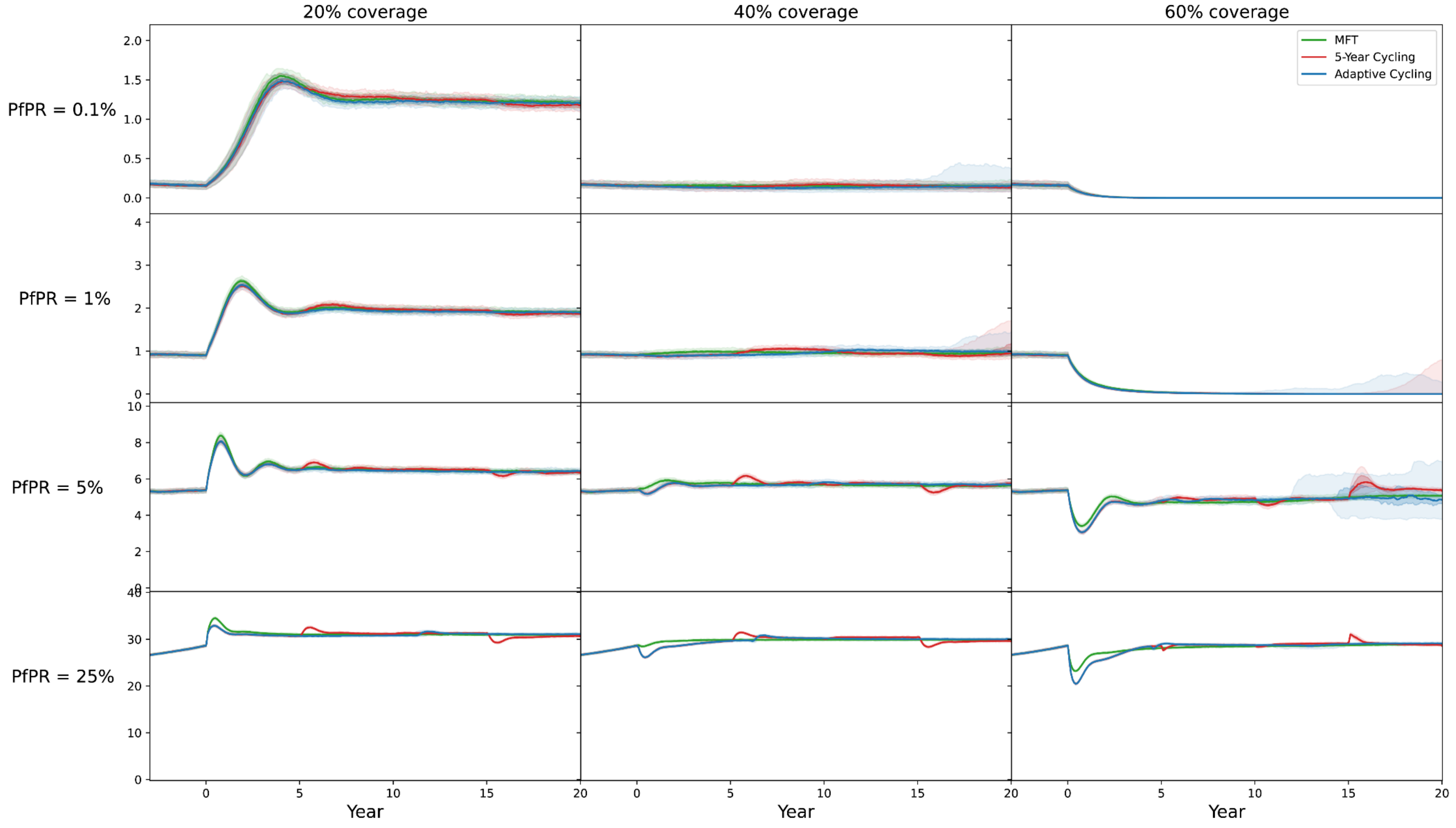


**Supplementary Figure 25.** Prevalence levels ($PfPR_{2-10}$) for all twelve prevalence-coverage scenarios evaluated in this analysis. Shaded area are 95% ranges. Visible shaded areas are blue (adaptive cycling) and red (5-year cycling).
